# Bid as a novel interacting partner of IRE1 differentially regulating its RNAse activity

**DOI:** 10.1101/572222

**Authors:** Samirul Bashir, Debnath Pal, Mariam Banday, Ozaira Qadri, Arif Bashir, Younis Mohammad Hazari, Nazia Hilal, Mohammad Altaf, Khalid Majid Fazili

**Affiliations:** Department of Biotechnology, University of Kashmir, Hazratbal J&K, India; Department of Computational and Data Science (CDS), Indian Institute of Science (IISc), Bengaluru, India; Institute of Biomedical Sciences, University of Chile, 8380453 Santiago, Chile

## Abstract

Unfolded protein response is a dynamic signalling pathway, which is involved in the maintenance of proteostasis and cellular homeostasis. IRE1, a transmembrane signalling protein represents the start point of a highly conserved UPR signalling cascade. IRE1 is endowed with kinase and endoribonuclease activities. The activation of the kinase domain of IRE1 by trans-autophosphorylation leads to the activation of its RNAse domain. RNAse domain performs atypical splicing of Xbp1 mRNA and degradation of mRNAs by an effector function known as Regulated IRE1 Dependent Decay (RIDD). The regulation of the distinctive nature of the IRE1 ribonuclease function is potentially mediated by a dynamic protein structure UPRosome that is an assembly of a huge number of proteins on IRE1. Here, we reported that Bid is a novel recruit to UPRosome, which directly interacts with the cytoplasmic domain of IRE1. Bid controls the auto-phosphorylation of IRE1 in a negative manner where Bid overexpression conditions displayed reduced phosphorylation levels of IRE1 and Bid knockdown cells showed slightly enhanced IRE1 phosphorylation. This effect was reciprocated with JNK, a downstream target of IRE1. Our Insilico analysis revealed that Bid binding to IRE1 dimer averts its structural flexibility and thereby preventing its trans-autophosphorylation activity. We found that the effect of Bid is specific to the IRE1 branch of UPR signalling and competitive in nature. The highlighting observation of the study was that Bid stimulated a differential activity of the IRE1 RNAse domain towards Xbp1 splicing and RIDD. These results together establish that Bid is a part of the UPRosome and modulates IRE1 in a way to differentially regulate its RNAse outputs.

## Introduction

Endoplasmic reticulum (ER) provides the proper microenvironment and the necessary tools to accurately fold proteins (Helenius *et al*., 2004). Owing to varied physiological and pathophysiological conditions, the protein folding capacity of the ER gets compromised, consequently leading to ER stress. To combat ER stress, cells have developed an ER to nucleus transcriptional network known as unfolded protein response (UPR). In mammalian cells, UPR is composed of three main signalling branches namely IRE1 (Inositol requiring enzyme 1 signalling branch), PERK (PKR-like ER kinase) and ATF6 (Activating transcription factor 6) (Schroder *et al*., 2005; Walter *et al*., 2011), while in yeast UPR is mainly operated through IRE1 branch. The net initial effect of UPR is to reduce ER stress by increasing the amount of ER-resident chaperones, ER luminal space, and other folding catalysts. However, if UPR fails to restore the homeostasis, it then initiates apoptosis in the cells (Jager *et al*., 2012).

The most conserved branch of the UPR signalling from yeast to humans is represented by the IRE1 signalling network. IRE1 is a type I transmembrane protein with dual enzyme activity (kinase and endoribonuclease) localized to the ER membrane. Under non-stressed conditions, IRE1 remains inactive through Bip binding but this inhibition is removed by induction of ER stress. Once activated, IRE1 performs splicing of Hac1 mRNA in yeast and Xbp1 mRNA in humans (Sidrauski *et al*., 1997; Gonzalez *et al.*, 1999; Calfon *et al.*, 2002). Additionally, RNAse domain of the IRE1 is involved in the degradation of a subset of mRNAs through a process referred to as regulated IRE1 Dependent Decay (RIDD) (Hollien *et al*., 2006; Hollien *et al*., 2009). The nature of Xbp1 splicing and RIDD activity is distinct. Xbp1 splicing is commonly linked to the cellular homeostasis whereas RIDD activity displays functional duality, where it can be cytoprotective or apoptotic depending on the stress conditions ER is facing. Though Xbp1 mRNA and RIDD substrates contain similar types of RNA architecture that are responsible for recognition and cleavage by IRE1 the discrepancy between these two activities is not clearly understood. Therefore, it is plausible to uncover the mechanistic details involved in IRE1 activation and function.

The dynamics of the IRE1 signalling is regulated by the association of several protein factors. IRE1 assembles in a huge protein complex designated as UPRosome (Hetz *et al*., 2009), where it is thought to act centrally and interact with several proteins, which can either activate/inactivate or connect IRE1 with other signalling networks. UPRosome as a multimeric protein complex has a great potential to explain the mechanistic details of IRE1 activation and its downstream effector functions particularly for understanding the dual nature of IRE1 activity. Several members of the Bcl-2 protein family have been found to interact with IRE1 and modulate its activity. Binding of Bak and Bax proteins to IRE1 stabilize the oligomer and maintain its activation. Additionally, the presence of Bak and Bax affected IRE1 mediated JNK activation (Hetz *et al*., 2006).

The members of the Bcl-2 protein family interact with each other; therefore it is possible that other members of the family may be recruited to UPRosome and regulates IRE1 dynamics. In this context, we screened interactions between IRE1 and the three proteins members from the Bcl-2 family namely Bid, Bik and Bcl-2. We found that Bid directly interacts with the cytoplasmic domain of IRE1 and modulates its activation. We observed that Bid specifically impacts IRE1 signalling and decreases the phosphorylation levels of IRE1 and its downstream target JNK. Interestingly, our results revealed that Bid differentially regulates two outputs of IRE1 RNAse activity, Xbp1 splicing, and RIDD. Bid repressed IRE1 can perform Xbp1 splicing but loses its RIDD activity. Our Insilco analysis predicted that this function of Bid is attributed to it’s binding at a crucial structural element of IRE1, which inhibits its transauto-phosphorylation activity and prevents the formation of higher oligomeric sate desired for the RIDD activity. We were able to show that Bid driven regulation of IRE1 activity is competitive in nature as overexpression of IRE1 leads to the restoration of RIDD activity. Together, our results added another regulatory element to UPRosome and are suggestive of an atypically novel function of Bid protein in the ER stress-induced IRE1 signalling pathway.

## Results

### 1. Structural analysis of IRE1 dimer

IRE1 structurally constitutes the sensory N-terminal domain towards the ER lumen, the C-terminal catalytic domain towards the cytosol, and a linker region connecting the two domains. IRE1 undergoes dimerization/oligomerization in the lumenal domain, which functionally activates the catalytic C-terminal domain. A quaternary structure of the IRE1 lumenal dimer was deduced from the crystal structure published earlier (Zhou *et al.*, 2006) (Figure 1A). The proposed quaternary structure with a large stable interface is formed through a beta-sheet spreading across the dimer subunits. Several mutation studies for residues 105, 123, 125 (Zhou *et al.*, 2006), and 109 (Liu *et al.*, 2003), corroborate the effect on dimer formation. There are, however, two problems with this form of the quaternary structure: (i) presence of a strong interface, which implies that a large activation and deactivation energy may be needed to alter the monomer-dimer equilibrium, which may not be adequate for a sensor protein, (ii) the end-to-end distance of the C-termini of lumenal domain between two monomers is 62.8 Å. For the lumenal domain to tether to the cytoplasmic domain across the membrane, a similar N-terminal end-to-end separation for cytoplasmic dimeric form of the protein is desirable. This can be found in the head-to-tail form of the dimeric proteins as shown in Figure 3A, where a 65.7 Å separates the dimer, which matches closely with the separation of C-terminal distance in the lumenal domain (Figure 1A). Also, there is a ~193 residue linker region that attaches the lumenal domain to the cytoplasmic domain with a 20 residue transmembrane segment located at positions 444-464. This means that there are roughly 76 and 98 residues on the lumenal and the cytoplasmic side that tethers the C-terminal domains, which must also dimerize to perform its biological functions. However, unless the 98-residue segment can form a stable structure, the lumenal---C-terminal domain tether is likely to remain floppy because of the large spatial gap between the two tethers.

**Figure1:**
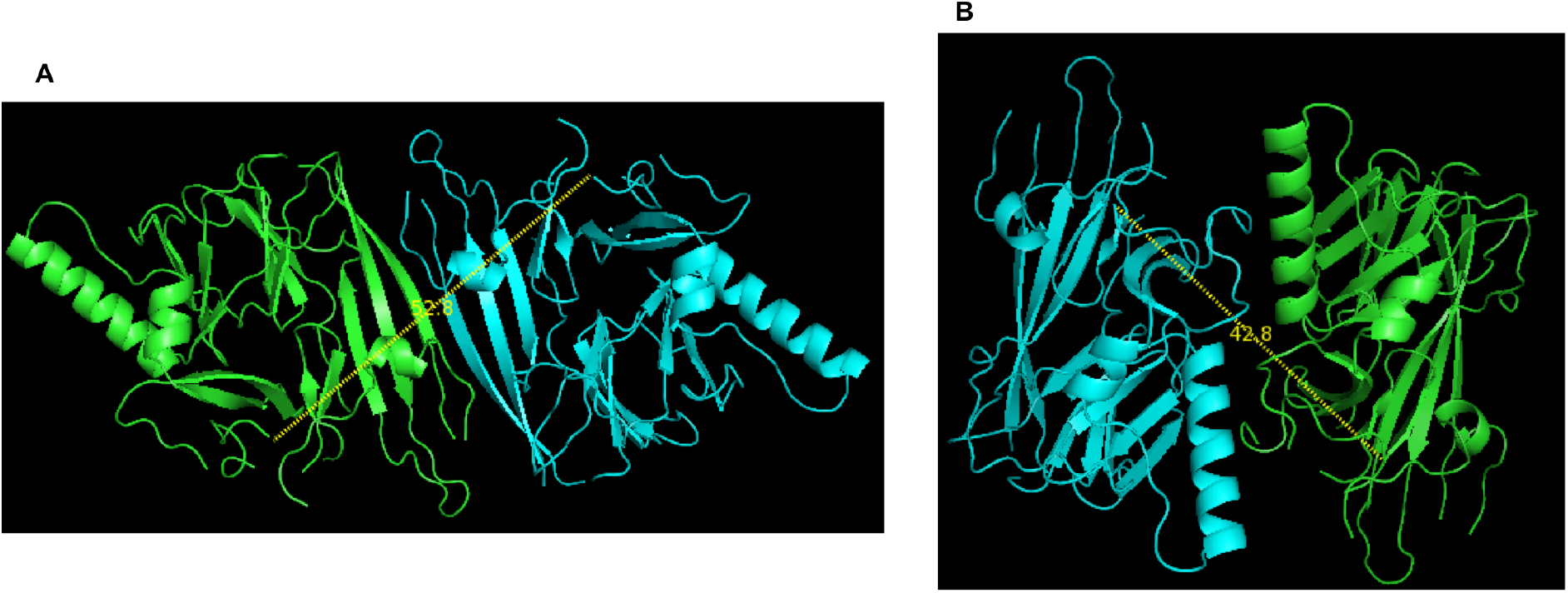
Quaternary structure form of the lumenal domain of IRE1 (PDB ID: 2HZ6). **A)** The more stable form of the quaternary structure of IRE1 with 62.8Å end-to-end distance between C-termini of monomers. **B)** Less stable form with 42.8 Å end-to-end distance between the C-termini of monomers. Distances between the C-terminal ends are depicted with a dashed line.

The structural perspective from the crystal structure data somehow does not fit well with the fragmented experimental data available, and therefore better strategies need to be developed to synchronize the data. To solve this structural discrepancy, here we are presenting an alternative quaternary structural model for the dimerization of the IRE1 lumenal domain (Figure 1B). The side-by-side orientation is obtained from the crystal lattice of the published structure and has been inferred as the potential dimer by our program. In this model, the C-termini of two monomers is separated by a shorter distance of 42.8Å. A corresponding C-terminal domain that matches has an N-terminal separation of 41.7 Å (Figure 3B). The C-terminal domain here is organized in a side-by-side orientation. Interestingly, this orientation has been suggested to be the correct dimerization state of the IRE1 C-terminal domain (Joshi et al., 2015). In our case, the lumenal domain dimer (Figure 1B) is a less stable form with 1130 Å of interface area, and −12.3 and 0.3 kcal/mol of theoretical interaction and dissociation free energy, respectively. In contrast, the other model depicts a more stable form of the quaternary structure of IRE1 (Figure 1A) with 1730 Å of interface area, and −15.2 and 7.2 kcal/mol of theoretical interaction and dissociation free energy, respectively. It means that in our model, the dimer has a reasonably large interface area and a low dissociation free energy indicating that it can easily undergo monomer-dimer transition. Also, an approximate 40Å separation between the luminal---C-terminal domain tethers is more likely to have a stable structure that stabilizes the full IRE1 protein in its functional form.

The activation of the catalytic activity of IRE1 requires its dimerization; however, the correct orientation of dimerization needed for activity is ambiguous because the cytoplasmic domain of protein showed head-tail dimerization in ADP bound form and side-by-side dimerization in the apo-form. This indicates that the interface dissociation energy is very low in both the dimeric states of the protein. The head-tail arrangement has an interface area of 1093 Å, but −1.6 kcal/mol of computed interface free energy contributed by 7 hydrogen bonds and 3 salt bridges. On the other hand, the side-by-side dimer has 1151 Å of interface area, has interaction free energy of +8.9 kcal/mol contributed by 33 hydrogen bonds and 32 salt bridges. It is obvious from the calculations that the side-by-side dimer is stabilized by entropic forces because theoretical free energy value is positive for a complex that has been experimentally determined. It also means that the stability of this complex will be tightly coupled to the immediate environment where it exists. On the other hand, the head-tail dimer is much more stable through enthalpic contributions and will need higher activation-deactivation energy to alter its dimerization state. The side-by-side dimer is more suitable to act as a sensor protein. These results collectively suggest that our quaternary structural model presents the desired conformation of IRE1 lumenal dimer that is in synchrony with the potential side-by-side orientation of cytoplasmic dimer.

### 2. Bid physically interacts with IRE1

The connection between Bcl-2 protein family and ER stress was demonstrated by an early report, where it was shown that Bax and Bak deficiencies conferred resistance to ER stress-mediated cell death (Pihán *et al*., 2017). The studies have also revealed that some of the Bcl-2 proteins like Bak, Bax, Puma, and Bim directly interact with IRE1 to regulate its functions (Hetz *et al*., 2006; Rodriguez *et al*., 2012). To get a fine-grained view of the role of Bcl-2 proteins in the Ire1 signalling pathway, it is essential to investigate whether the other members of the family have any role in UPR signalling pathway. To this end, interactions between IRE1 and Bcl-2 family members such as Bid, Bik, and Bcl-2 were screened by immunoprecipitation assay. Full length Ire1 (untagged) was co-expressed with GST-tagged Bid, or Bik, or Bcl-2 in HEK293T cells. Cells either transfected with IRE1 and GST-tagged vector, or un-transfected were used as a negative control to demonstrate the specificity of the assay. This was followed by tunicamycin (Tm) treatment to induce ER stress. For the immunoprecipitation assay, a 6hr time point was chosen because it was speculated that Bcl-2 proteins might be recruited to UPRosome at the later stages of ER stress. The IRE1 immunoprecipitated samples using the anti-IRE1 antibody were run on SDS-PAGE and probed for the presence of Bcl-2 proteins by using an anti-GST antibody. The results from this experiment revealed that IRE1 interacts with Bid, while no interactions were found with Bik and Bcl-2 (Figure 2A). The interaction between IRE1 and Bid was amplified by increasing the doses of plasmid DNAs as depicted by the enhanced signals on the blot (Figure 2B). The presence of IRE1 in pull-down samples was confirmed by probing the blots with anti-IRE1 antibody (Figure 2A and B). The interaction was further verified by reverse immunoprecipitation where pull-down was carried by anti-GST antibody followed by western blot with anti-IRE1 antibody (Figure 3C). From these results, we concluded that among the Bcl-2 proteins, only Bid forms a complex with IRE1.

**Figure2:**
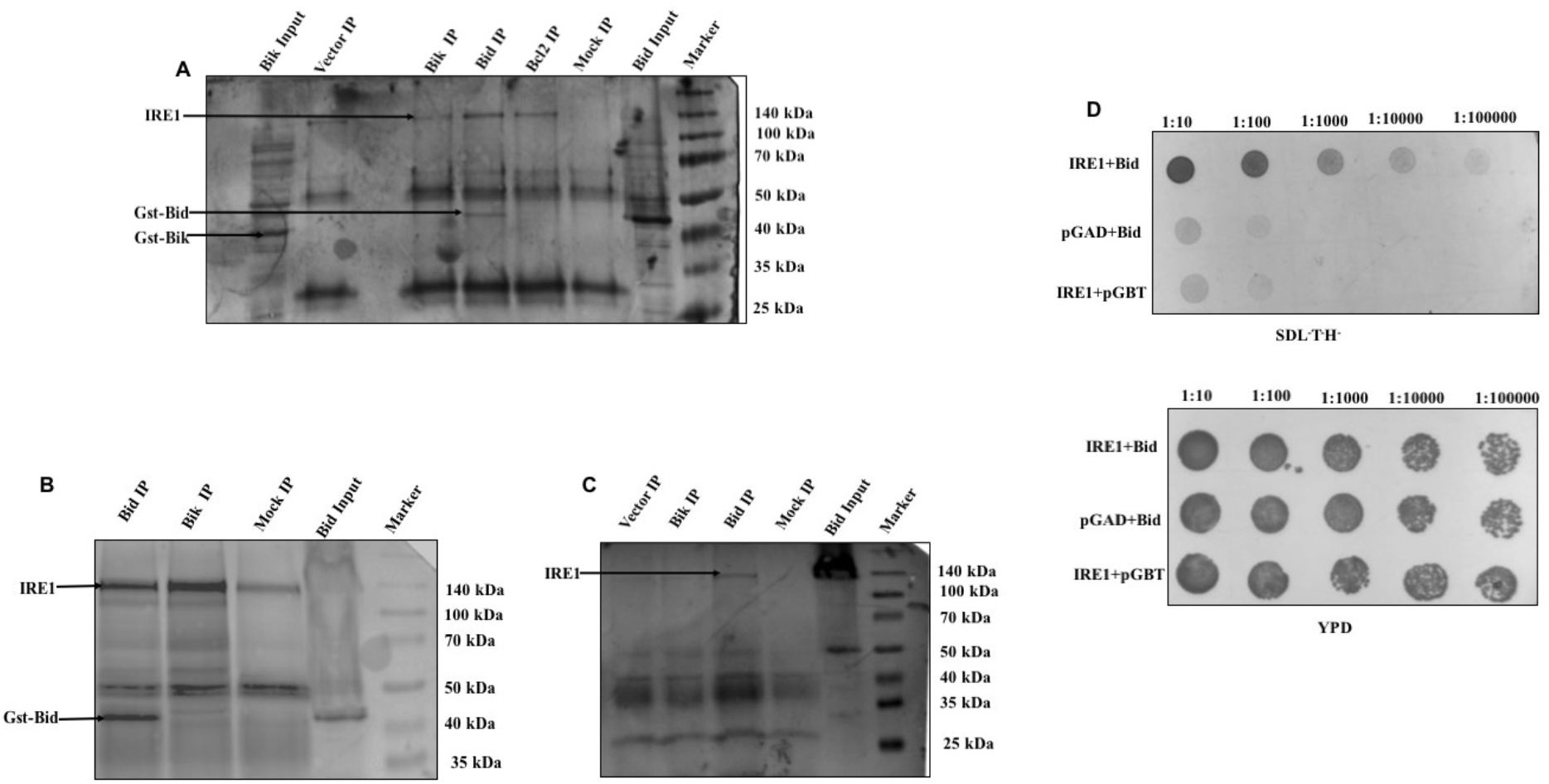
Bid physically interacts with IRE1. **A)** HEK293-T cells were co-transfected with full-length IRE1 and Gst-Bid, IRE1 and Gst-Bik, IRE1 and Gst-Bcl2 and IRE1 and Gst-Vector. Cells were treated 6μM tunicamycin for 6hr. Immunoprecipitation was performed with the anti-IRE1 antibody. The presence of Bcl2 proteins in the IRE1 complex was detected by western blot analysis using an anti-Gst antibody. Bid IP, Bik IP, BCl-2 IP, and Vector IP corresponds to immuno-pulldown experiments for Gst-Bid, Gst-Bik, Gst-Bcl2, and Gst-vector (pEBG) co-expressed with Ire1 respectively. Mock IP represents HEK293-T cells without transfection. Marker represents the protein molecular mass indicator. **B)** Repeated Immuno-precipitation experiment for Bid and Bik. Blots were also probed with an anti-IRE1 antibody to check the presence of IRE1 in pulldown samples. **C)** Immuno-precipitation experiment for Bid and Bik with Gst-antibody. **D)** pGAD-Ire1 + pGBT-Bid represents colonies obtained from the co-transfection of pGAD-IRE1 and pGBT-Bid plasmids. pGAD424 + pGBT-Bid shows colonies obtained from co-transfection of pGAD424 and pGBT-Bid plasmids. pGAD-IRE1 + pGBT9 represents colonies obtained from the co-transfection of pGAD-IRE1 and pGBT9 plasmids. Dilution spotting on SDL^−^T^−^H^−^ drop out media to select for positive interactors (Upper panel). Lower panel shows dilution spotting on YPD rich media. Dilutions were made up to 10^−5^.

**Figure3:**
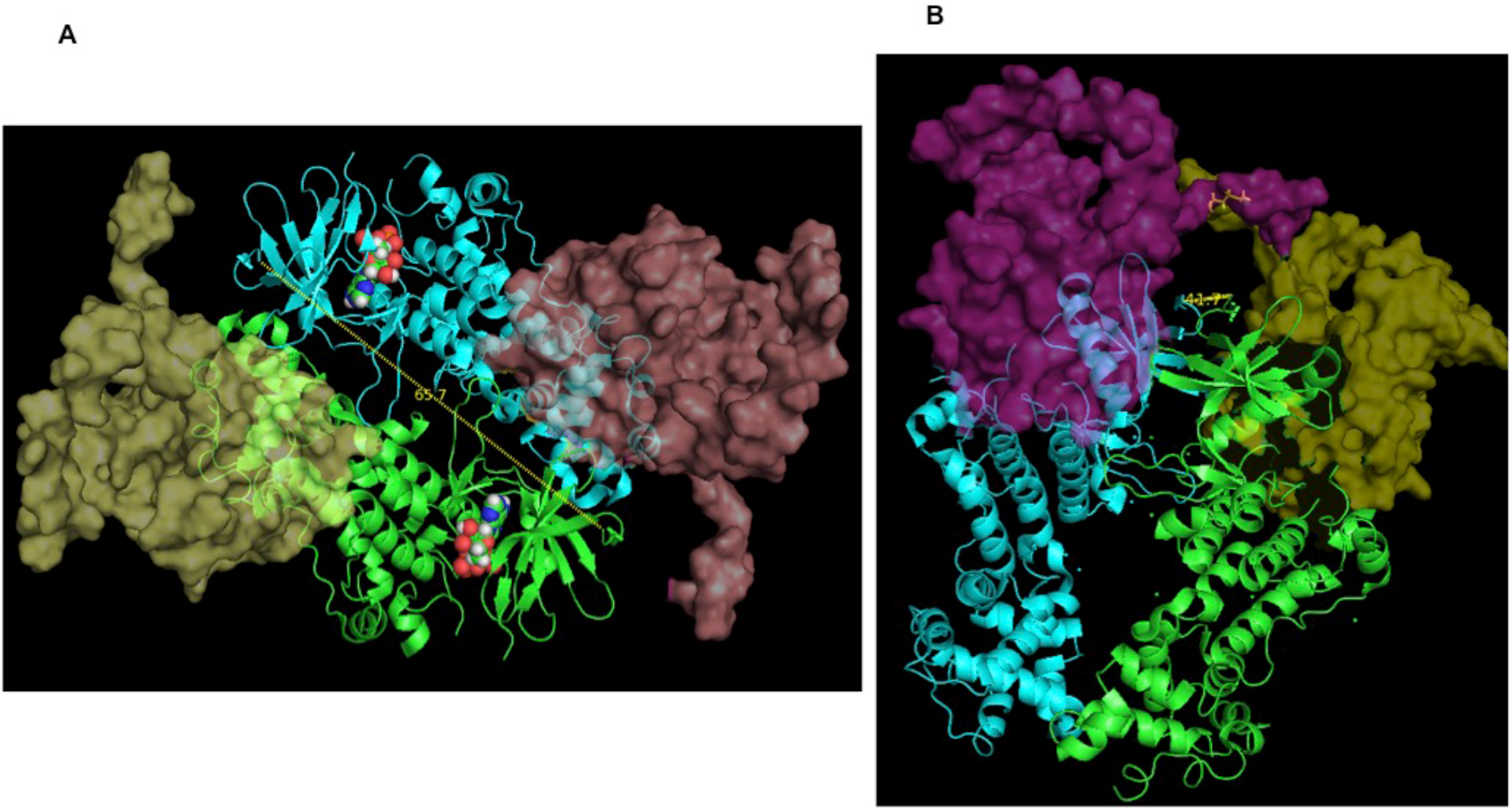
Bid interacts with the IRE1 cytoplasmic domain. **A)** A cartoon diagram of dimeric IRE1 bound in a head-tail form (PDB ID: 4YZD). One subunit of the monomeric BID is docked on to the N-terminal domain of IRE1 and shown as transparent-surface models. The end-to-end N-terminal distance of the dimers is also shown in a dashed line and 65.7 Å distance marked on it. The bound ADP is shown in CPK model. **B)** Cartoon diagram showing the side-by-side dimer quaternary structure of IRE1 in apo form; i.e., no bound ligand (PDB ID: 4Z7G; chains cyan and green) docked by two subunits of monomeric protein BID (PDB ID: 2BID; surface colors pink and yellow). The docking poses are obtained from the first ranked hits from a scan utilizing ZDOCK software. Coordinates from 4U6R PDB ID was taken for docking because it had the most completely determined IRE1 structure in monomeric form.

To characterize the nature of the interaction between IRE1 and Bid, yeast two-hybrid assay was carried out. IRE1 was translationally fused with the GAL4 activation domain (GAL4AD) and Bid was fused with the GAL4 DNA binding domain (GAL4BD). AH109 strain of *Saccharomyces cerevisiae* that harbors a chromosomally integrated His (Histidine) reporter gene was used for the experiment. AH109 cells co-transfected with IRE1 and Bid grew well in histidine dropout media compared to controls (Figure 3D). This implies that in IRE1+Bid co-transfected colonies, the GAL4 transcription factor was successfully reconstituted by the interaction between IRE1 and Bid, which leads to the expression of His-reporter gene and allowed growth on His drop out media. The viability of the cells was demonstrated by spotting them on yeast rich media (Figure 3D). Therefore, we conclude that Bid shows a direct physical interaction with IRE1.

### 3. Bid interacts with the cytoplasmic domain of IRE1

From the immunoprecipitation and the yeast two-hybrid assays, it was observed that Bid directly interacts with IRE1. Since IRE1 is a huge protein with different domains, it is imperative to identify the region of Bid docking on IRE1. For docking, we selected the cytoplasmic domain of IRE1 because Bid is primarily localized towards cytoplasm and rationally should interact with the IRE1 cytosolic domain. By using ZDOCK software, a predictive docking experiment was run to find the Bid docking regions of IRE1 protein. For our study, 4U6R (Crystal structure of IRE1 cytoplasmic domain complex with a ligand) was chosen as a reference protein because it existed in monomeric form and has the most completely determined crystal structure at the highest resolution to date. It may be noted that the Pro51-Gln81 stretch of Bid is of irregular structure. The N-terminal segment Gly1 to Glu16 is also of irregular structure. Therefore, although we got hits from the ZDOCK for all varieties of poses, in such cases where the irregular structural segment makes a contact, they were deemed as ambiguous results and therefore discarded. The rank 1 complex from the software output, which had a score of 808.658 was taken for investigation. The docked positions can be seen from Figure 3. These figures were created after the superposition of docked 4U6R onto the respective IRE1 models from 4YZD (crystal structure of phosphorylated IRE1 in complex with ADP-Mg2+) and 4Z7G (crystal structure of IRE1 in the apo form). As can be seen from the figures, the contact of Bid is with the N-terminal region of the cytoplasmic domain of IRE1 protein and contacting the sheet-turn segment located between positions 565-595 and 627-635. These results reveal that BID interacts with IRE1 through its cytoplasmic domain.

### 4. Bid negatively regulates the phosphorylation of IRE1 and its downstream target JNK

The UPRosome represents a molecular platform, where association and dissociation of proteins dynamically regulate IRE1. Although, UPRosome is composed of a large number of proteins presence or absence of each protein has a significant effect on IRE1 transauto-phosphorylation activity (Hetz *et al.*, 2009; Chen *et al.*, 2013). In this context, we wanted to analyze the effect of Bid on IRE1 phosphorylation status. For this, the Bid gain of function and loss of function studies were carried out in HEK293T cells. Cells were transiently transfected with Bid or vector, or untransfected followed by induction of ER stress using 6μM tunicamycin for 6hr. For Bid depleted conditions, cells were transfected with Bid siRNA or negative control siRNA, or untransfected followed by tunicamycin treatment. siRNA knockdown conditions were standardized for Bid for 24 to 48hr and selected 42hr time point for subsequent knockdown experiments (Figure S1). Immunoblotting was performed to detect the levels of phopho-IRE1 by using an anti-IRE1 (phospho S724) antibody. Our results revealed that Bid overexpression drastically decreased the levels of phospho-IRE1 compared to controls (Figure 4A). Interestingly, our results also revealed that the levels of IRE1 phosphorylation increase in Bid knockdown conditions (Figure 4B). Total IRE1 levels remained unchanged (Figure S2). These results, therefore, concluded that Bid negatively regulates IRE1 phosphorylation.

**Figure4:**
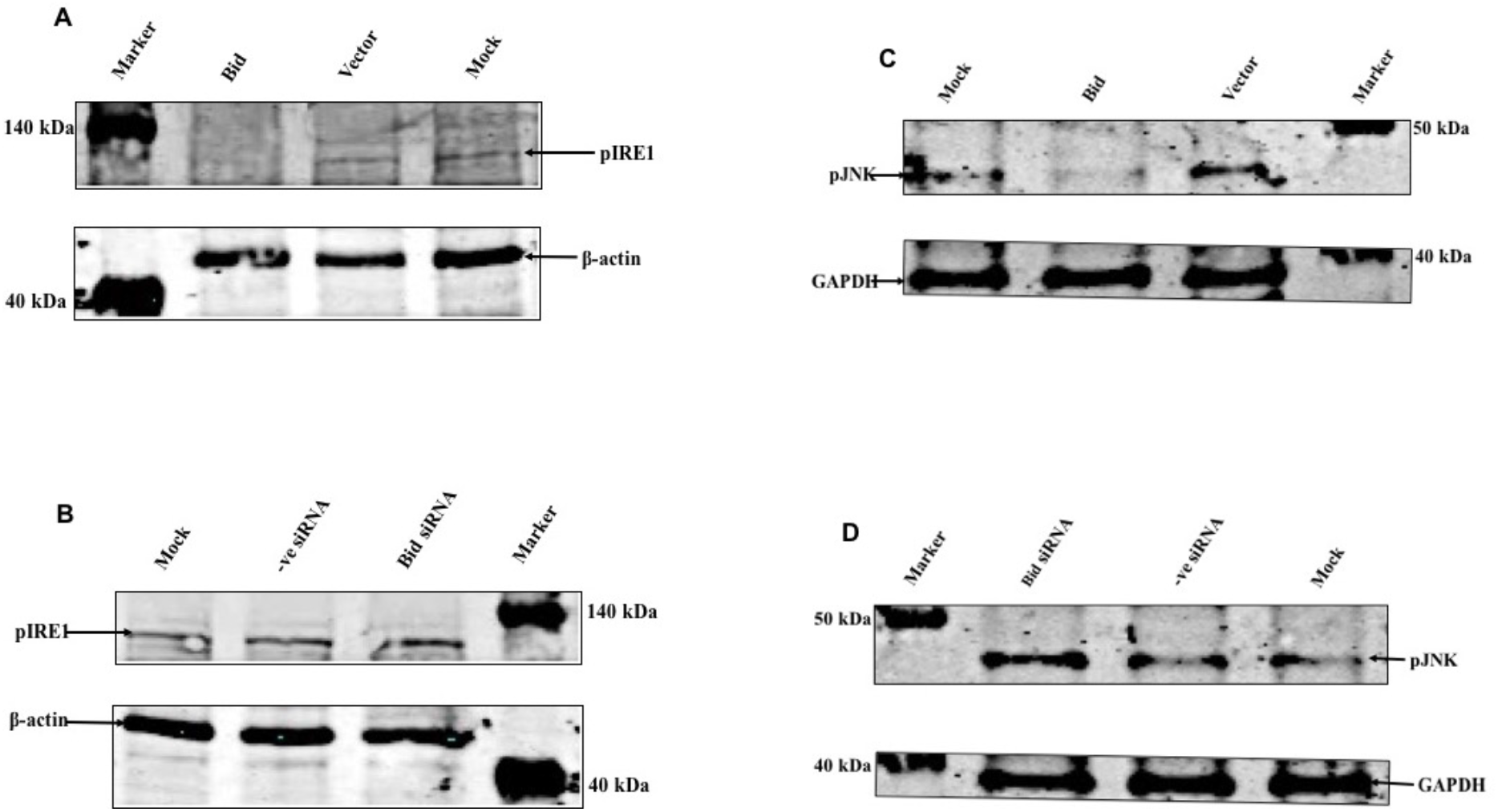
Bid negatively regulates phosphorylation of IRE1 and JNK. **A)** Analysis of IRE1 phosphorylation in Bid overexpressing conditions. HEK293-T cells were transiently transfected for Bid, Vector and Mock. After 42hr cells were stimulated with 6μM tunicamycin for 6hr. Levels of pIRE1 were checked by probing with the anti-IRE1 (phosphoS724) antibody. **B)** Analysis of IRE1 phosphorylation in Bid knockdown conditions. HEK293-T cells were transfected with Bid siRNA and -ve siRNA followed by treatment with 6μM tunicamycin for 6hr. **C)** HEK293-T cells were transiently transfected for Bid, Vector and Mock. After 42hr cells were stimulated with 6μM tunicamycin for 6hr. **D)** HEK293T cells were transfected with Bid siRNA and -ve siRNA followed by treatment with 6μM tunicamycin for 6hr. Western blot analysis was performed to check the levels of pJNK by probing with Phospho-JNK Rabbit mAb. β-Actin and GAPDH represent the endogenous controls for pIRE1 and pJNK respectively. Marker represents a Pre-Stained Protein Ladder, Bid (overexpressed Bid), Vector (overexpressed vector), Mock (no transfection control), -ve siRNA (mission siRNA fluorescent universal negative control #1, Cyanine 3) and Bid siRNA (Mission esiRNA human BID).

Several studies have revealed that activation of IRE1 upon ER stress is associated with the activation of JNK (Urano *et al*., 2000; Yoneda *et al*., 2001). Since our experiments revealed negative regulation of IRE1 phosphorylation driven by Bid, it was plausible to investigate the effect of Bid on phosphorylation of JNK. Similar sets of experiments were designed to check the effect of Bid on JNK phosphorylation. Inconsistent with IRE1 phosphorylation status, it was observed that there is a decline in JNK phosphorylation upon Bid overexpression (Figure 4C). Similarly, JNK phosphorylation was enhanced in Bid depleted conditions (Figure 4D). Together these results suggest that Bid modulates JNK phosphorylation in a similar fashion to IRE1.

### 5. Bid specifically impacts IRE1 arm of UPR

UPR signalling is mediated by three branches in metazoans, IRE1, PERK, and ATF6. It is thought that in metazoan grafting of IRE1 gave rise to the evolution of PERK, where the luminal domain of IRE1 fused with the eIF2α kinase domain. Also, the secondary structure analysis revealed a similarity between IRE1 and PERK. This observation was further substantiated by a study where it was found that the lumenal domain of yeast IRE1 can be functionally substituted for PERK lumenal domain from other species (Bertolotti *et al.*, 2000; Liu *et al.*, 2002). It might be possible that PERK activation might follow a trend similar to IRE1. Therefore, the effect of Bid on the PERK arm was evaluated. Once activated PERK catalyzes the phosphorylation of eIF2α that puts a block on global translation. However, under these conditions, the translational activation of a transcription factor (ATF4) is enhanced due to the presence of an upstream ORF (uORF). So, we detected the levels of phospho-eIF2α and ATF4 to analyze the effect on the PERK arm. Bid overexpression and knockdown conditions were adapted for the analysis. Western blotting was performed to detect the levels of peIF2α by probing with an anti-peIF2α antibody. Similarly, the levels of ATF4 were detected by western blot probing with an anti-ATF4 antibody. It was found that the levels of eIF2α, as well as ATF4, remain unchanged in both Bid overexpression and knockdown conditions (Figure 5A and B). Therefore, we concluded that the PERK arm is not influenced by Bid.

**Figure5:**
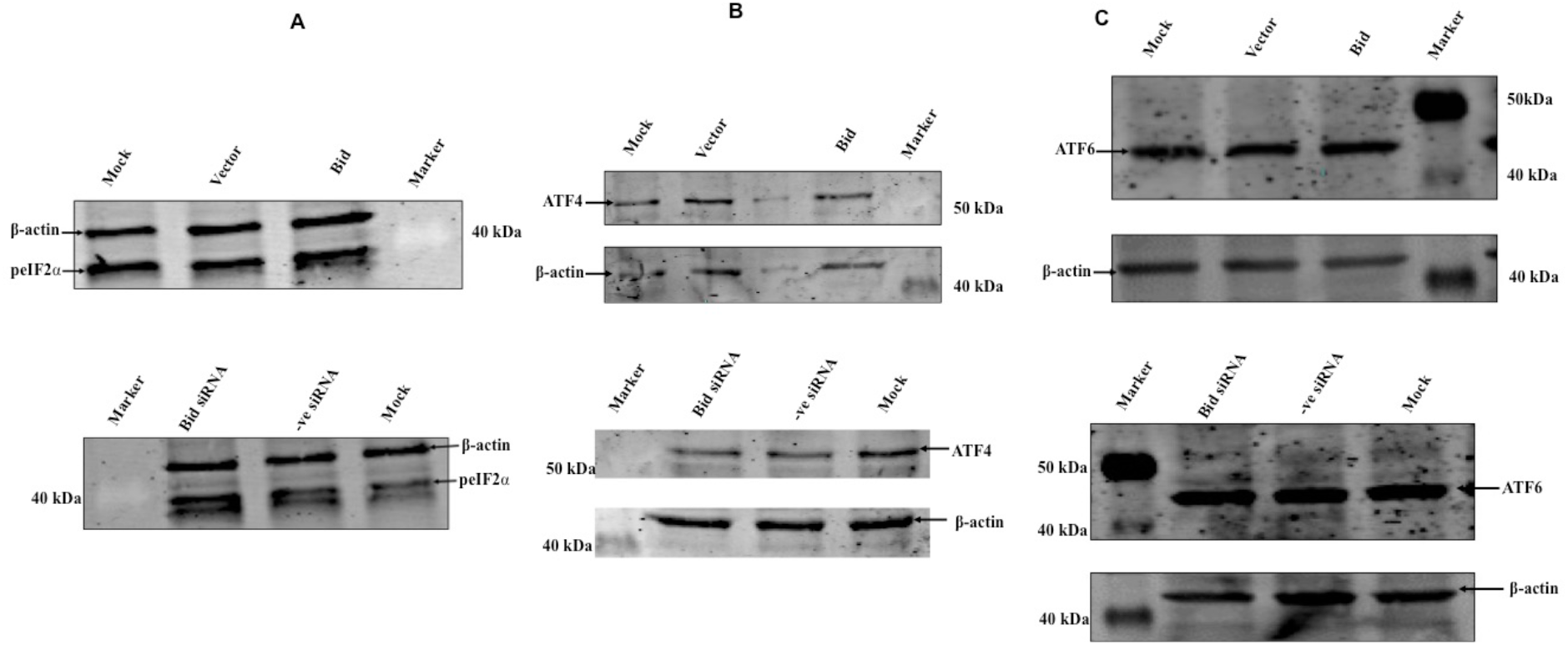
Effect of Bid on PERK and ATF arms of UPR. HEK293T cells were transiently transfected with Gst-tagged Bid, Gst-tagged Vector and Mock (no transfection). After 42hr cells were stimulated with 6μM tunicamycin for 6hr. HEK293T cells were transfected with Bid siRNA and -ve siRNA followed by treatment with 6μM tunicamycin for 6hr. **A, B, C)** Western blot was performed to check the levels of phosphor-eIF2α **(A)**, ATF4 **(B)**, and ATF6 **(C)** by probing with Phospho-eIF2α antibody, anti-ATF4 antibody, and ATF6 antibody respectively. β-Actin represents the endogenous controls. Marker represents a Pre-Stained Protein Ladder, Bid-Gst (overexpressed Bid), Vector-Gst (overexpressed vector), Mock (no transfection control), -ve siRNA (mission siRNA fluorescent universal negative control #1, Cyanine 3) and Bid siRNA (Mission esiRNA human BID).

ATF6 branch of UPR follows a different mechanism for activation. There is no similarity between the lumenal domains of ATF6 and IRE1. Also, ATF6 is present in an oligomeric state under normal conditions, unlike IRE1 and PERK. However, binding of Bip and its release is common for both IRE1 and ATF6. To evaluate that the effect of Bid is not a general effect rather a specific effect, we analyzed the effect on the ATF6 branch of UPR. The activation of ATF6 is detected by the presence of its cleaved active transcriptional factor. Both gains of function and loss of function conditions were used for the analysis. Immunoblotting was performed to detect the levels of cleaved ATF6 by probing with the anti-ATF6 antibody (Figure 5C). We observed that there is no difference in the levels of cleaved ATF6 under all experimental sets. We concluded that like PERK arm; the ATF6 arm of the UPR remains unaffected. This leads to the conclusion that Bid specifically regulates the IRE1 branch of UPR.

### 6. Bid differentially modulates IRE1-RNAse outputs

The comprehensively documented function of IRE1 is performed by its RNAse domain, which catalyzes the splicing of Xbp1 mRNA by excising a 26-nucleotide intron. IRE1 phosphorylation is important for the catalysis of Xbp1 splicing and it has been shown that for successful Xbp1 splicing, phosphorylation of kinase activation loop becomes indispensable. This was further supported by an observation where unmitigated splicing of Xbp1 was detected in a phosphomimetic mutant of yeast IRE1. In our experiments, a decline in IRE1 phosphorylation levels in the presence of overexpressed Bid protein was observed. To evaluate the effect of Bid on Xbp1 splicing, levels of the spliced Xbp1 protein was analyzed. Surprisingly, it was found that spliced Xbp1 protein remained unchanged in both overexpression and knockdown experimental conditions (Figure 6A and B) through the levels of pIRE1 changed in response to Bid. The status of Xbp1 splicing was also checked by semi-quantitative RT-PCR. Because of the presence of 26-nucleotide intron, unspliced Xbp1 (Xbp1u) runs slower than the spliced mRNA product. Here, also we didn’t find any change in Xbp1 splicing status (Figure 6C). This result was further confirmed by analyzing a downstream target of Xbp1, Bip. Levels of Bip protein did not show any significant changes in response to Bid (Figure 6D). These results pointed towards the fact that Xbp1 splicing operates even under low IRE1 phosphorylation states.

**Figure6:**
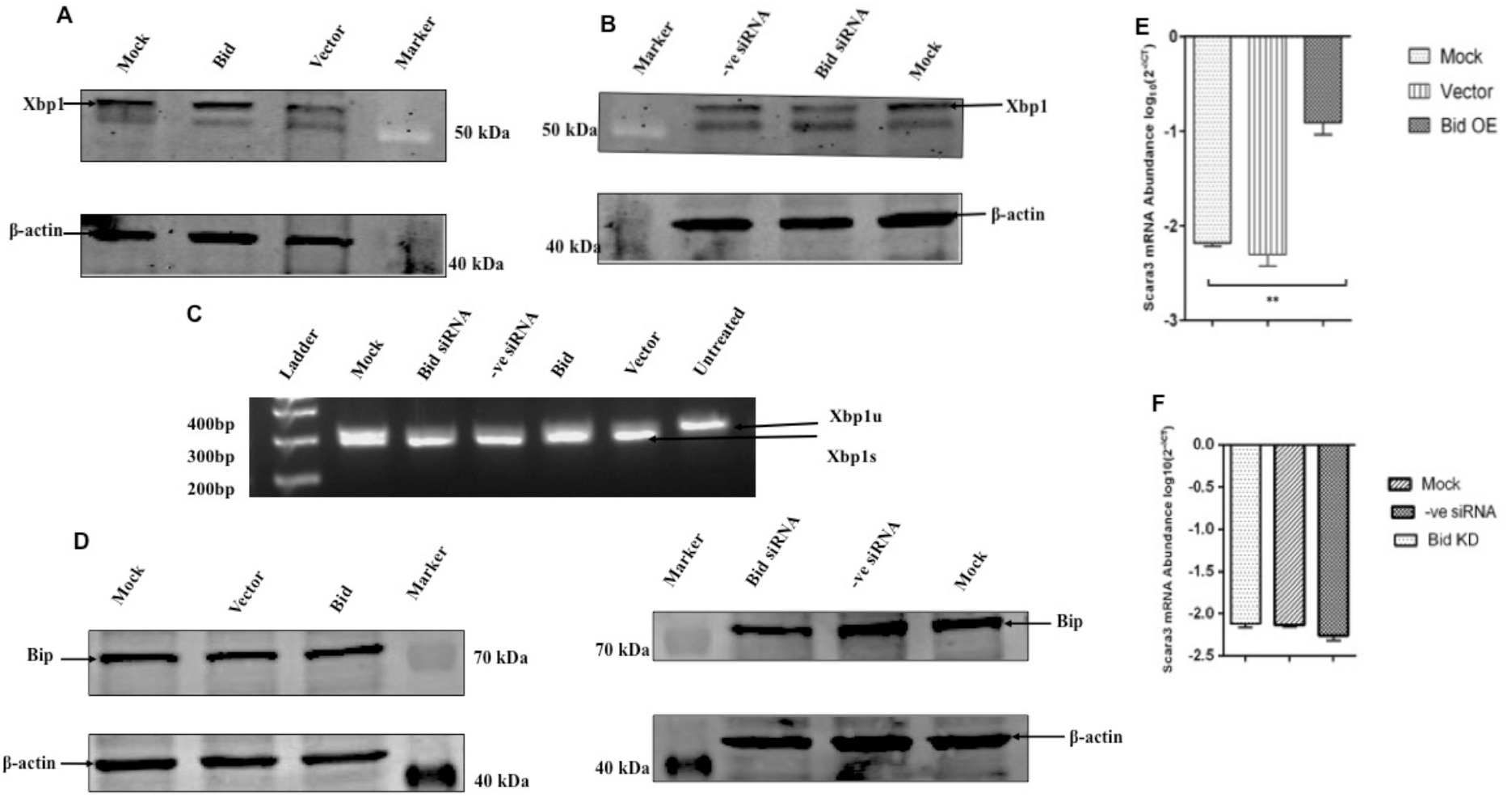
Bid differentially modulates IRE1 RNAse outputs. **A)** HEK293-T cells were transiently transfected for Bid, Vector and Mock. After 42hr induction with 6μM tunicamycin was given for 6hr. Western blot was done to check the levels of spliced Xbp1 by probing with the anti-Xbp1 antibody. **B)** Bid siRNA was transfected in HEK293-T cells along with a negative control siRNA. 6hr treatment with 6μM tunicamycin was given after 42hr of transfection. Western blot was done to check the levels of spliced Xbp1 by probing with the anti-Xbp1 antibody. **C)** XBP-1 splicing detected by semi-quantitative RT-PCR. PCR products were separated on 2% agarose gel. The upper band defines unspliced Xbp1 (Xbp1u), where the lower band is spliced Xbp1 (Xbp1s). **D)** Levels of Bip protein was checked by probing with Anti-Bip antibody, β-Actin represents the endogenous control. **E and F)** Abundance of Scara3 mRNA was analyzed by real-time PCR. Real-time PCR quantitation was done by plotting log_10_ of 2^−ΔCT^ and represent mean SD values. **E** and **F** represent overexpression and knockdown respectively. Statistical analysis was performed using One-way ANOVA: **: p < 0.0016.

The RNAse domain of IRE1 is also involved in the degradation of mRNAs preferably localized to ER through RIDD. RIDD activity is also dependent on the phosphorylation status of IRE1. For evaluating the effect of Bid on RIDD activity, we analyzed the levels of scara3 (a known RIDD target) by real-time PCR analysis. Interestingly, it was found that scara3 mRNA levels are elevated in Bid overexpression compared to controls, which illustrates the reduced RIDD activity in Bid overexpressing conditions (Figure 6E). We also observed slightly improved RIDD activity upon Bid knockdown (Figure 6F). These results demonstrated that RIDD activity has severely defected in Bid overexpression state that is inconsistent with a decrease in IRE1 phosphorylation levels. Our results are in corroboration with a study, where they showed that modulating IRE1 phosphorylation/oligomerization displays differential Xbp1 splicing and RIDD outputs (Han *et al.*, 2009). Taken together these results, we suggest that Bid differentially regulates IRE1 RNAse activity.

### 7. Bid-mediated IRE1 regulation is competitive in nature

It is well established that Bip keeps IRE1 in an inactive state under non-stress conditions but this inhibition is disrupted by the induction of ER stress. Increased levels of unfolded proteins in ER titrate out Bip from IRE1 and leads to its activation (Bertolotti *et al.*, 2000; Zhou *et al.*, 2006; Oikawa *et al.*, 2009). Similarly, we propose that concentration-dependent events are operating in the case of Bid mediated IRE1 regulation. To test this hypothesis, Ire1 was co-expressed with either Bid or Vector in HEK293T cells. Mock (untransfected cells) was used as a negative control and all the samples were treated with 6μM tunicamycin for 6hr. Western blot analysis was done to detect levels of pIRE1 by probing with anti-IRE1 (phospho S724) antibody. It was observed that the levels of pIRE1 were almost similar in IRE1-Bid co-expression conditions and controls (Figure 7A). This result unveiled that IRE1 overexpression could release the Bid repression. Next, the effect on IRE1 downstream targets Xbp1 splicing and RIDD activity was analyzed. Levels of spliced Xbp1 and scara3 (RIDD target) were detected by real-time PCR. We observed increased Xbp1 splicing in cells co-expressing Ire1 and Bid, and Ire1 and vector compared to Mock (Figure 7B). This was expected as increased levels of IRE1 enhances Xbp1 splicing. Strikingly, we observed enhanced degradation of scara3 in cells when Ire1 and Bid were co-expressed (Figure 7C). These results are in contrast to our earlier results, where we observed that Bid overexpression inhibits Scara3 degradation; this implies that Ire1 overexpression can rescue RIDD activity. These results point towards a mechanistic model, where Bid and IRE1 work competitively to regulate the RNAse activity of IRE1.

**Figure7:**
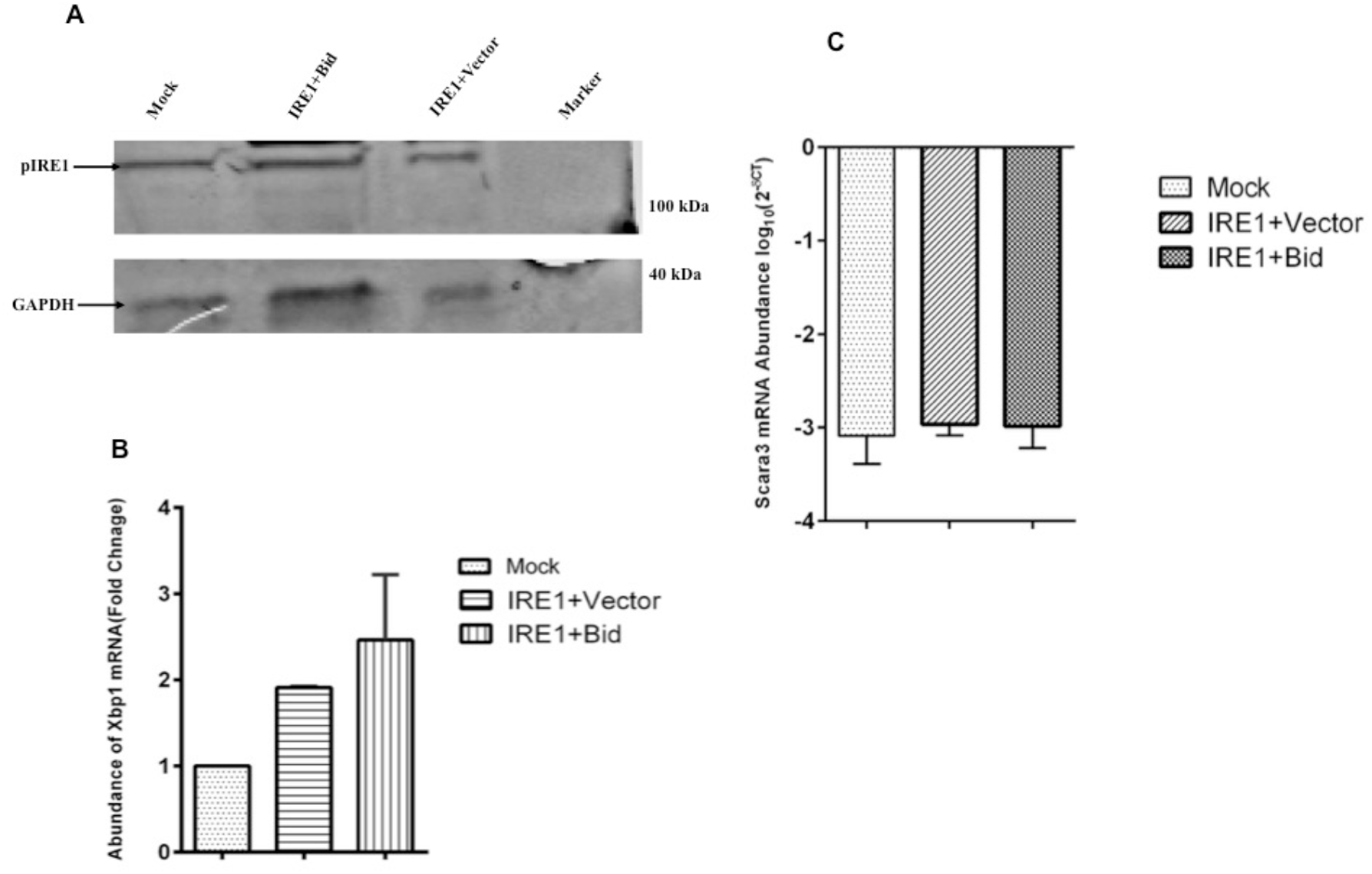
Bid competitively inhibit IRE1 activation. Ire1 was co-expressed with Gst-tagged Bid and Gst-tagged Vector independently in HEK293T cells. Mock was kept as no transfection control. Cells were then treated with 6μM tunicamycin for 6hr. **A)** Western blot analysis was done to detect the levels of pIRE1 using the anti-IRE1 (phospho S724) antibody. GAPDH was used as an endogenous control. **B and C**) RNA was extracted and real-time PCR analysis of Xbp1 spliced mRNA **(B)** and Scara3 mRNA **(C)**. Xbp1s were plotted as fold change. Scara3 mRNA levels were plotted as log10 values. Statistical analysis was performed using One-way ANOVA and represent as mean SD.

### 8. Mechanism of Bid-mediated regulation of IRE1 trans-autophosphorylation

The observations from docking results suggested that Bid docked structure in no way blocks the interface as found in the dimers in the PDB 4YZD and 4Z7G. It also does not block the ATP binding pocket to obstruct access to the ligand. Bid also does not bind to the catalytic domain to obstruct any catalysis site as per our best docking result. Therefore, why does the interaction of Bid with the IRE1 C-terminal domain obstruct trans-autophosphorylation and its catalytic activity? To answer these questions, we did the multiple sequence alignment for IRE1 protein, in which secondary structures obtained from three different crystal structures of IRE1 were aligned. Labels 1 and 2 correspond to dimers from the apo form of IRE1 (PDB ID 4Z7G), A and B correspond to a dimer of IRE1 in ADP bound form (PDB ID 4YZD), and label a and b correspond to monomers from 4U6R superposed onto 4YZD dimer (Figure 8). An overall look at the alignment reveals very limited regions of conserved regions (yellow color), indicating the limited existence of a full consensus in secondary structure, which is the expected observation since the primary structure is 99% identical in these proteins. Interestingly, we see that the edge of the regular secondary structures has altered in each protein, and in some cases, there is a transition from strand to helix secondary structure and vice versa. The most significant alterations of our interest lie in and around the ATP binding pocket since that is the reaction center for the autophosphorylation step. The first key observation is that the activation loop is missing in the crystal structure 4YZD and 4Z7G (Figure 9). This indicates that the activation loop is a dynamically flexible region of the protein. The ATP binding site is comprised of residues 577-585, 643-645, and 690-693. These regions are part of beta-strands that alter conformation on binding/dimerization (Figure 8). These strands are part of the sheet that connects to the Bid-docking site. When Bid docks, the structural alterations are obstructed, and the ATP binding is impaired. The extent of structural alteration and consequence movement of the beta-sheet and the associated turns can be observed from Figure 9. A corroboration of this mechanism can be obtained from the mutation at site 599 (Tirasophon *et al.*, 1998; Iwawaki *et al.*, 2001) that is known to lead to loss of autophosphorylation and of endoribonuclease activity, and this site is close to the ATP pocket and the beta-sheets of interest. Since there is structural alteration, the complementary surfaces cannot form as required for specific binding and therefore the dimerization event would also be affected. These observations reveal that Bid binding to IRE1 can thereby obstruct its trans-autophosphorylation activity and dimerization events for both side-by-side and head-tail cases.

**Figure8:**
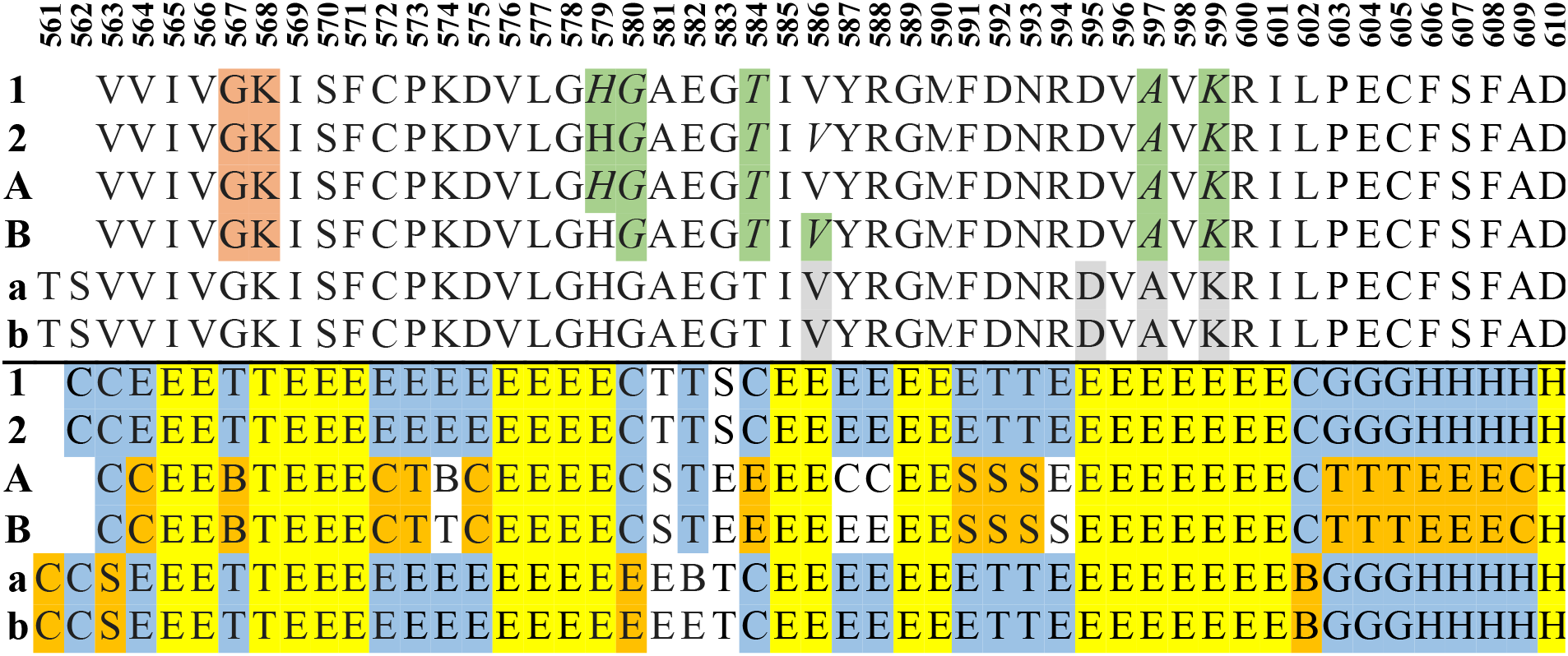
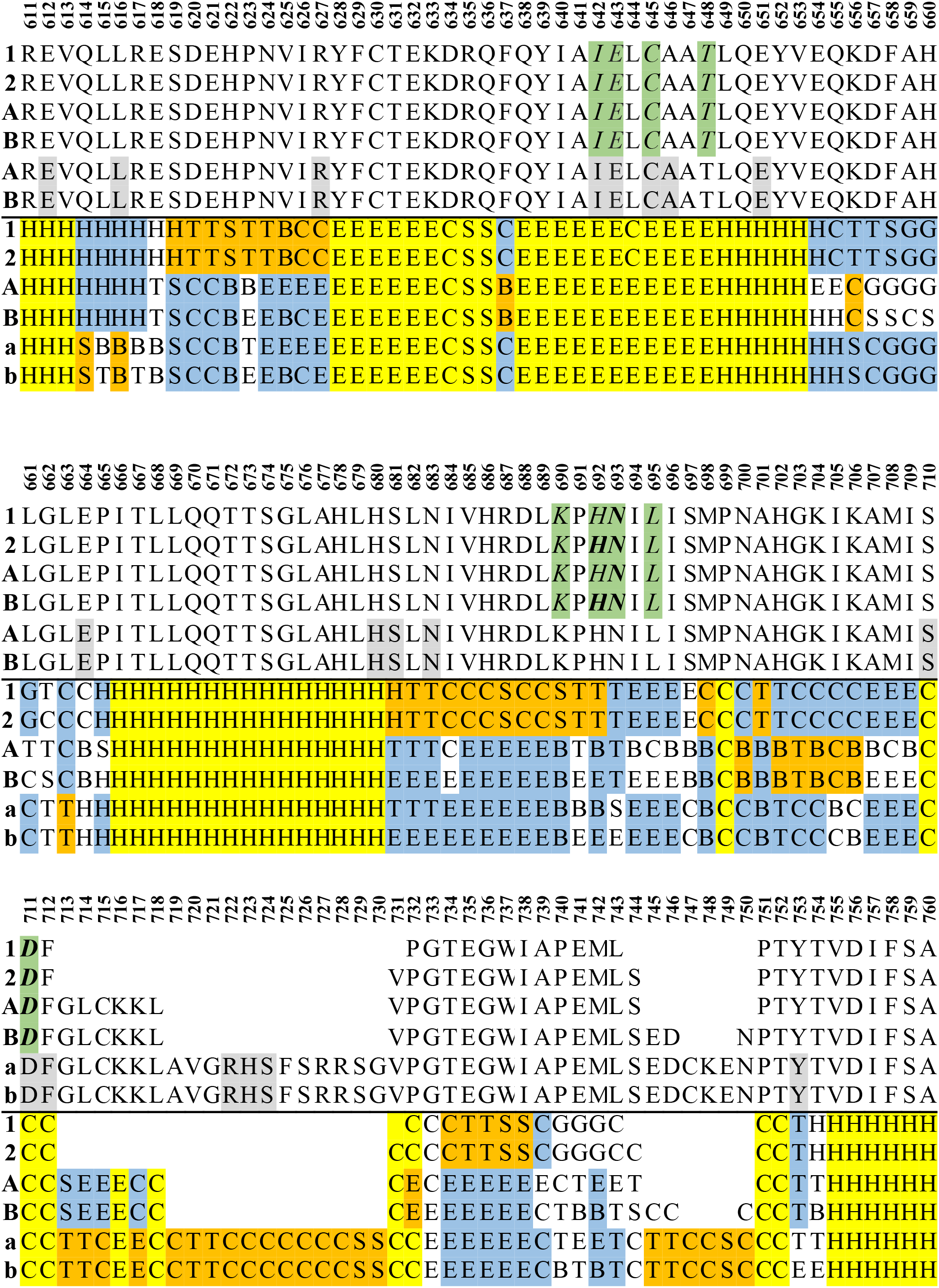
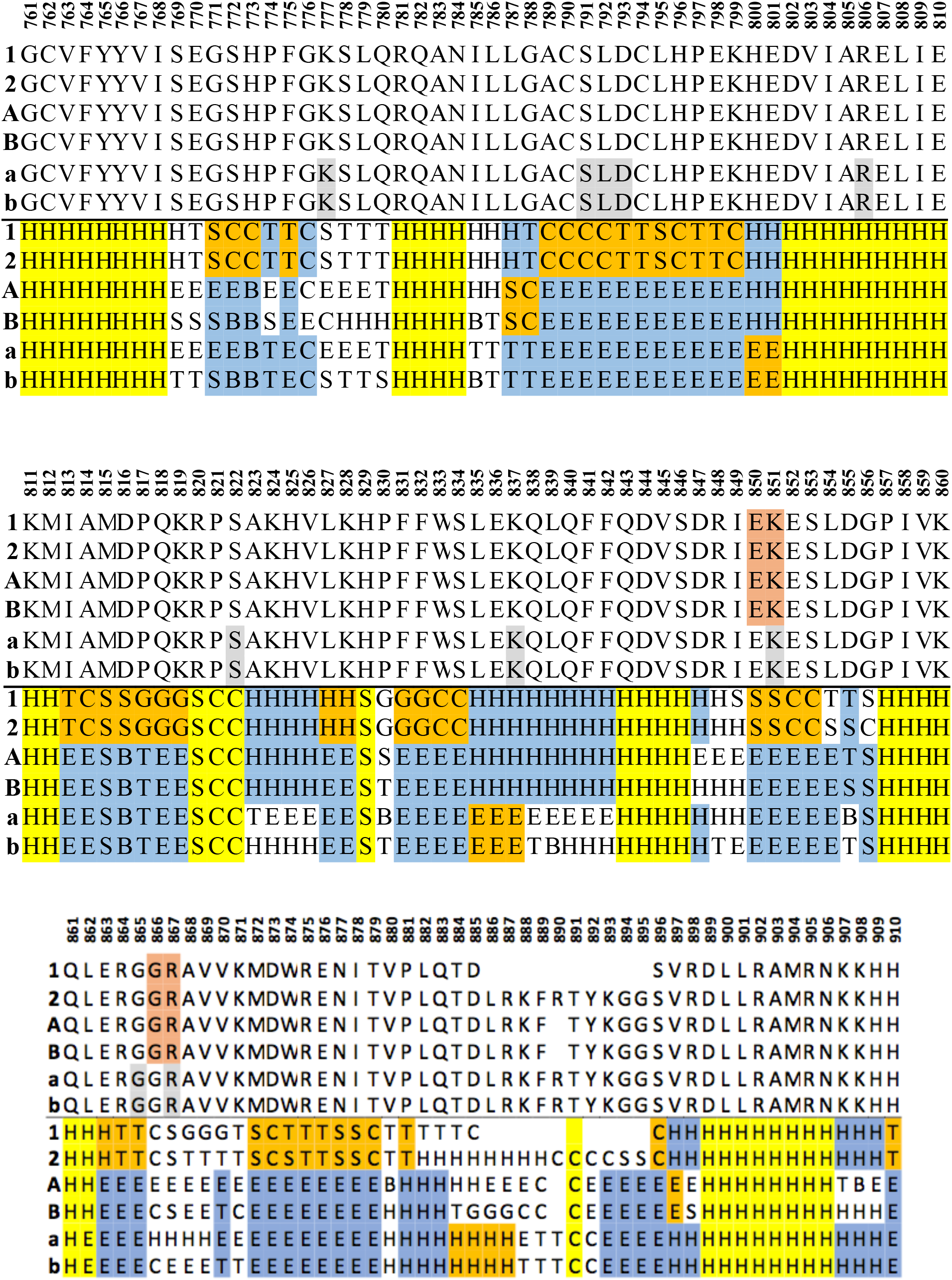
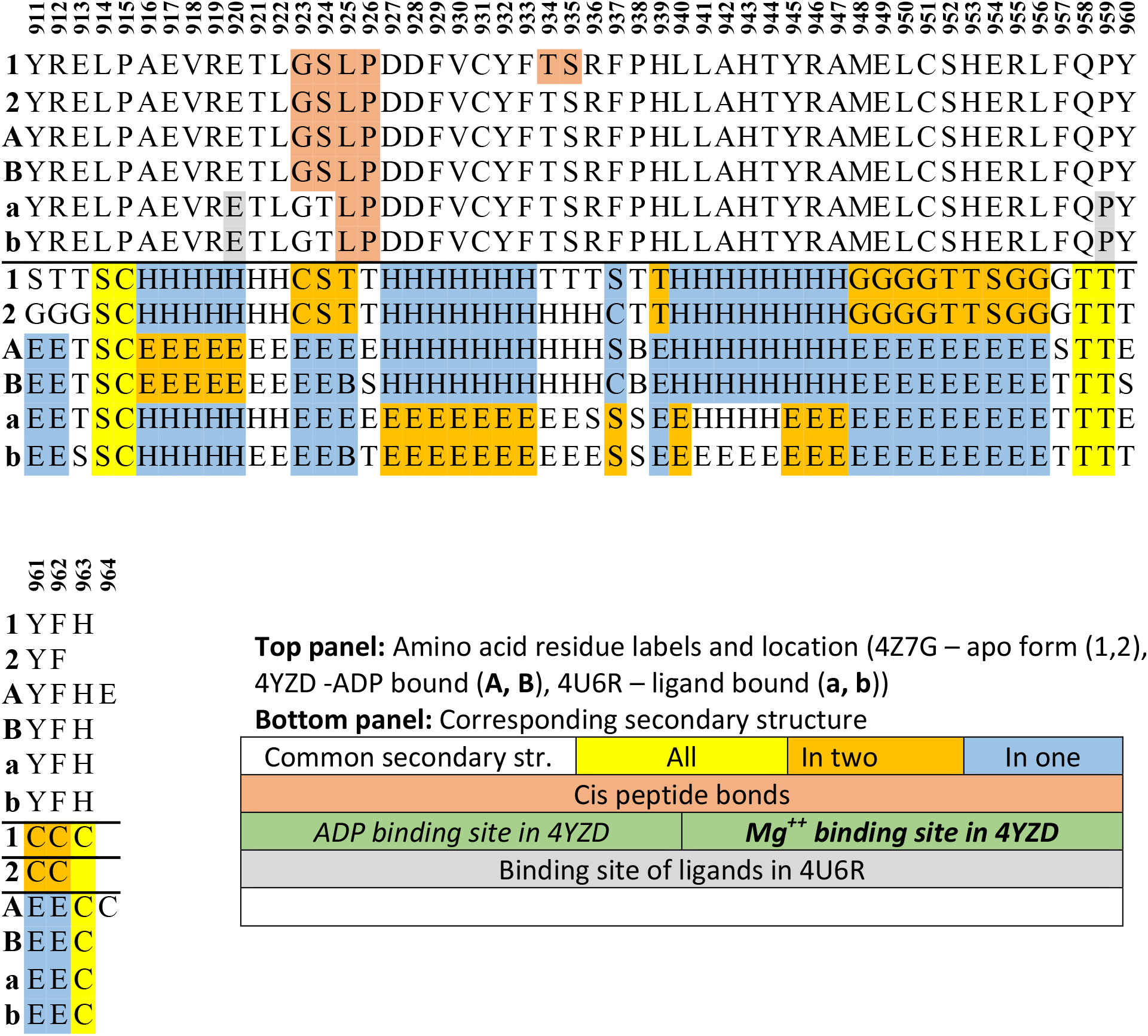
Structure-based multiple sequence alignment of amino acids and secondary structures present in three different protein structure files of IRE1 C-terminal domain proteins, 4Z7G, 4YZD, and 4U6R. The former two exist in dimeric form, and the latter in monomeric form. The secondary structure for two sequences of 4U6R was obtained after superposition onto the dimer of 4YZD. Segments, where the amino acid coordinates are absent in the protein structure files, are given as blank. Note that the apo- and the ADP-bound form of the proteins have six cis peptide bonds indicating large strain in the polypeptide backbone. Note the large alteration in the secondary structure in the ligand-bound form of the proteins, where the structure undergoes significant alteration. The ligand molecule for 4U6R is 3E4 (N- {4-[3-{2-[(Trans-4 Aminocyclohexyl) Amino] Pyrimidin-4-Yl} Pyridin-2-Yl-Oxy]-3-Methylnapthalen-1-Yl}-2 Chlorobenzene-sulphonamide), a large ligand that almost fills up the ATP binding pocket similar to what ATP would have done.

**Figure9:**
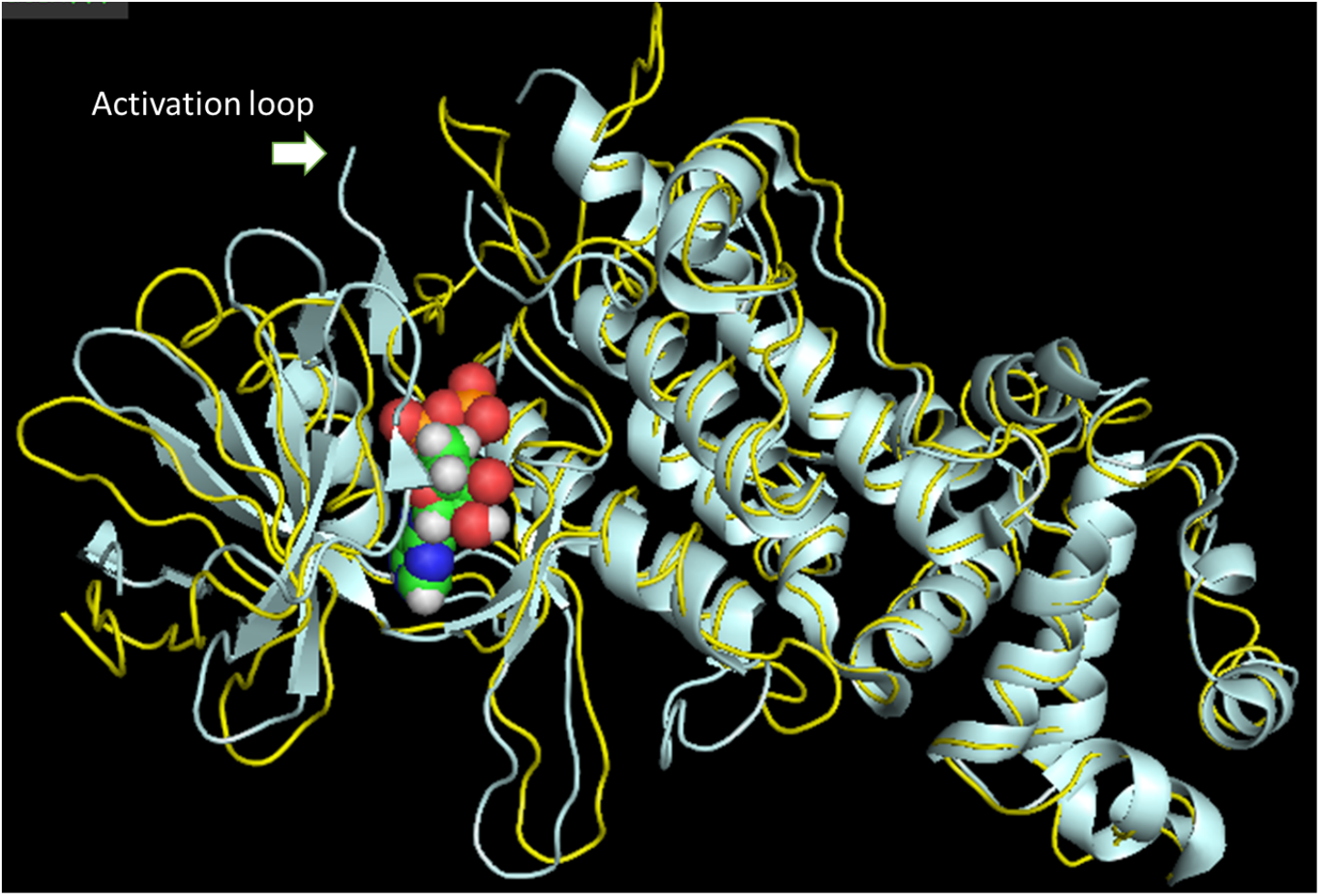
Superimposed IRE1: Cartoon diagram of one subunit of IRE1 in ADP bound form (PDB ID: 4YZD) superposed on 3E4 ligand-bound form (PDB ID: 4U6R). The activation loop is missing from the ADP bound structure and is marked by an arrow. The bound ADP is shown in CPK model.

## Discussion

IRE1 branch of the UPR signalling network represents the highly conserved pathway among the three branches and it provides a major platform for deciding the fate of the cells under ER stress. IRE1 harbors dual enzyme activity, kinase, and endoribonuclease. Activated IRE1 catalyze the non-canonical splicing of Hac1 mRNA in yeast and Xbp1 mRNA in humans (Sidrauski *et al.*, 1997; Gonzalez *et al.*, 1999; Calfon *et al.*, 2002) and degrades a subset of mRNAs localized to ER through RIDD (Hollien *et al.*, 2006; Hollien *et al.*, 2009). The activation of IRE1 depends on the presence of positive and negative regulators. UPRosome was identified as a huge protein complex assembled at IRE1 that regulates the dynamics of UPR signalling pathway. Protein factors recruited to UPRosome either activates or deactivates IRE1 in a stress-dependent manner (Hetz *et al.*, 2009; Woehlbier *et al.*, 2011). One of the interesting facts about the UPRosome is that factors recruited to it act atypically and perform their functions differently from their originally designated functions. Bak and Bax proteins have been found associated in a complex with IRE1, thereby activate its downstream target Xbp1 and promote cell survival. To better understand and unravel the role of UPRosome in regulating the activity of IRE1, which form the central core component of the UPRosome, we investigated the role of other Bcl-2 family members like Bid, Bik, and Bcl-2. We found that Bid is a novel interacting partner of IRE1 that modulates its activation and downstream signalling.

Bid is a BH3 only protein that upon cleavage of the C-terminal fragment (tBid) binds to Bak and Bax and promotes permeabilization of OMM and cytochrome C release (Billen *et al.*, 2009). Like Bak and Bax, we found that Bid also interacts with IRE1 and modulates its activation. The interaction is direct in nature as revealed by yeast two-hybrid assay. Here, we provided evidence for negative regulation of IRE1 phosphorylation driven by Bid. We observed a transition in phosphorylation levels of IRE1 between different states of Bid. Overexpression conditions of Bid decreased IRE1 phosphorylation drastically, and improved IRE1 phosphorylation levels were observed in Bid depleted conditions. These two contrasting conditions substantiated the fact that Bid has a negative effect on IRE1 phosphorylation. This regulatory function of Bid was found to be specific to the IRE1 branch of UPR because we did not find any perturbation in the other two arms of UPR (ATF6 and PERK) in Bid overexpression and knockdown conditions. Also, JNK phosphorylation levels were found decreased upon Bid overexpression and slightly increased in Bid depleted conditions. JNK activation is linked to the phosphorylation of through the IRE1-TRAF2 module. Consistent with IRE1 phosphorylation, JNK phosphorylation levels follow a similar trend. IRE1-JNK line has been linked to promoting cell death in response to chronic ER stress, thus places Bid in a position to promote cell survival by disrupting IRE1-JNK activation. These results might partially explain the role of Bid in cell proliferation (Bai *et al.*, 2005; Yin *et al.*, 2006). However, we need to further validate these results in a model organism and investigate the cell survival role of Bid in response to ER stress.

One of the interesting findings of our results revealed that Bid differentially activates IRE1 RNase activity. It was found that overexpression of Bid severely inhibited the degradation of RIDD target scara3. This effect was reversed in Bid knockdown conditions, where we observed slightly enhanced RIDD activity. In both the conditions, levels of spliced Xbp1 mRNA remained unchanged. This differential RNase activity is possibly linked to the phosphorylation state of IRE1. Our results are in corroboration with the previous studies where binding of 1NM-PP1 (an ATP analog) to kinase-dead mutant I642G Ire1 rescued Xbp1 splicing partially but failed to activate the RIDD (Papa *et al.*, 2003; Han *et al.*, 2009). These studies also lead to the observations that ATP-competitive inhibitors can allosterically activate RNAse domain of IRE1. In the case of wild type IRE1, APY29 (type I kinase inhibitor) was found to decrease trans-autophosphorylation. This kind of inhibition limited RIDD activity but allowed efficient splicing Xbp1 (Han *et al.*, 2009). IRE1 phosphorylation is directly associated with the formation of a higher-order oligomeric structure. Averting the formation of oligomeric IRE1 would alter its ribonuclease function (Ghosh *et al.*, 2014). It is thought that Xbp1 represents the most preferred substrate for the IRE1 RNAse domain that is why even in lower-order oligomeric structure it is being spliced. However, for RIDD activity higher oligomeric state of IRE1 is inevitable. Similarly, we speculate that Bid mediated suppression of IRE1 phosphorylation is responsible for preventing the generation of higher-order IRE1 oligomer and allows only Xbp1 splicing to occur. This presumption is supported by our Insilco data. We observed that Bid binds to the cytoplasmic domain of IRE1 at a very crucial structural element, which is around the ATP-binding pocket. Our multiple sequence alignment results revealed that ATP-binding pocket is highly altered in secondary structure between different crystal structures of cytoplasmic IRE1 domain indicating that this region is highly flexible and desired for dimerization/oligomerization. So, when Bid binds to a sheet-turn segment, it stabilizes the structure and prohibits structural flexibility, which is otherwise important for ATP binding and trans-autophosphorylation. Additionally, Bid docking mediated structural stability influences the formation of complementary surfaces for dimerization/oligomerization. Bid binds IRE1 between regions 565-595 and 627-635 that might have other functional importance apart from influencing structural flexibility. A couple of studies have suggested an important role of Tyr628 in regulating IRE1 autophosphorylation activity. Mutating Tyr628 to Tyr628Ala or Tyr628Leu completely inhibited the autophosphorylation activity of IRE1. This study also revealed that mutation of Tyr628 to Tyr628Phe enhanced autophosphorylation activity. These results together suggested a mechanism for the activation of RNAse domain through a positional rearrangement from buried to surface-exposed Tyr628 side chain (Joshi *et al.*, 2015; Yang *et al.*, 2016). Extrapolating these results to our study, we suggest that Bid binding to this region would halt this structural rearrangement and limit the availability of Tyr628 at the surface and thereby impair autophosphorylation and RNAse activity. We concluded that Bid binding impacts the formation of required dimeric/oligomeric fully active state and therefore differentially controls the IRE1-RNAse activity. We also suggest that Bid competitively traps IRE1 in a dimeric or a lower oligomeric state, thereby preventing RIDD activation. This statement is also supported by one of our finding that overexpression of Ire1 in presence of Bid rescued IRE1 phosphorylation and RIDD activity completely. We propose a model, where the presence of Bid inhibits higher-order oligomeric IRE1 structure formation and thus keeps RIDD under check, but fully supports Xbp1 splicing (Figure 10). However, to better understand this scenario, we have to monitor the effect on IRE1 oligomerization in the presence of the Bid.

**Figure10:**
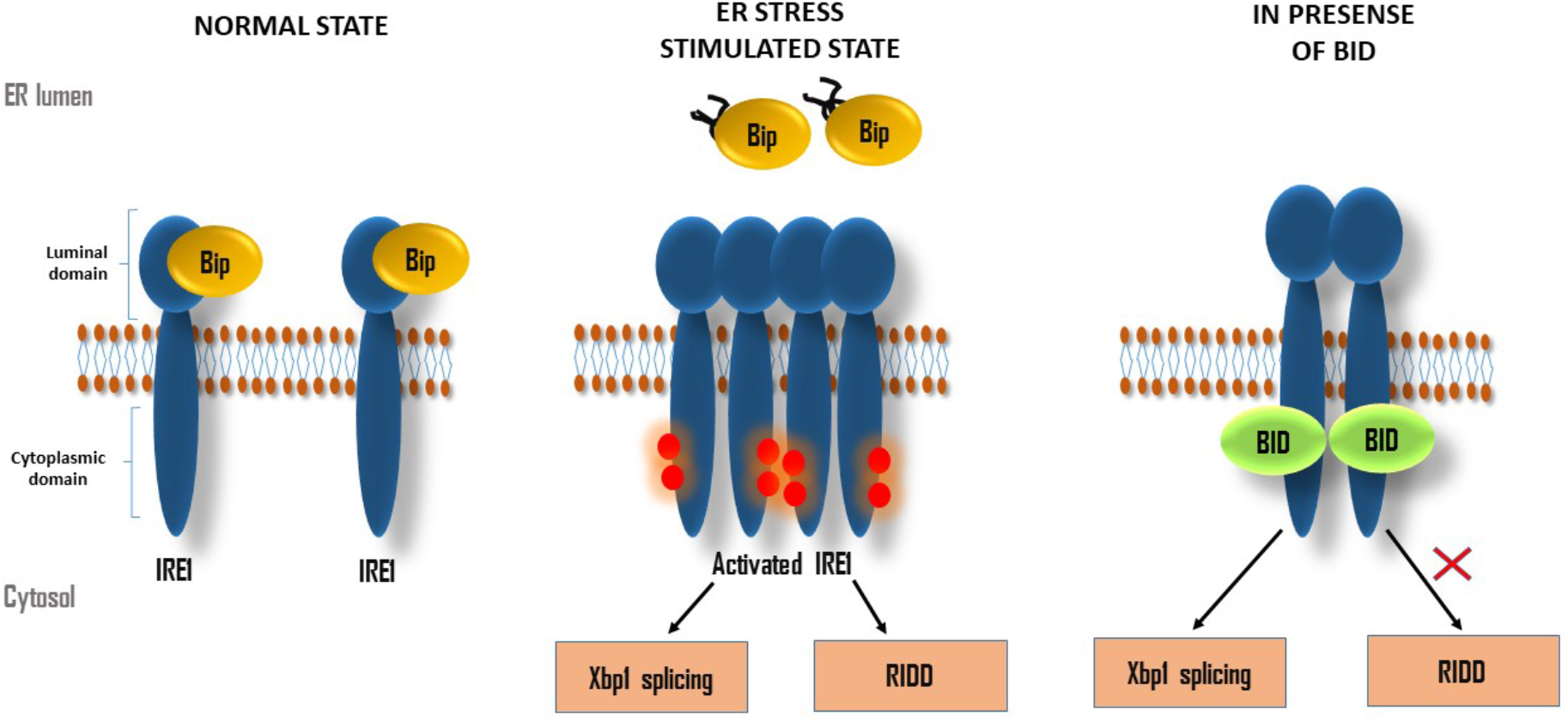
Model for Bid mediated regulation of IRE1 activity. Under normal conditions, IRE1 remains inactive due to Bip binding. However, Bip release induced by ER stress leads to IRE1 activation, which is followed by splicing of Xbp1 and degradation of mRNAs through RIDD. Binding of Bid to IRE1 cytoplasmic domain halts higher-order oligomer formation and thus allows only Xbp1 splicing to occur, while RIDD activity is reduced.

Does our study reveal any further observation that might contribute to understanding Bid interaction and autophosphorylation? Ser724 and Thr734 along with Ser726 and Ser729 residues constitute a part of the kinase activation loop, which is important for RNAse activity. In particular, Ser724 and Ser726 phosphorylation are pivotal for the activity of RNAse domain of IRE1. Residues 719-730, 745-750 are not visible in the 4YZD structure with ADP, 713-730, 744-750 are not visible in 4 Z7G in the apo form, while they are visible in 4U6R structure with 3E4. Loops 713-730, 744-750 are contiguous in space, and the former is part of the activation loop. However, 4YZD and 4Z7G structures have six cis peptide bonds. The situation is exactly opposite in 4U6R, where the activation loop is visible and fixed in a precise conformation, and this structure has only one cis peptide bond and that too far away from the ATP binding pocket. This suggests that the protein exists in multiple states where it is strained in some points but relaxed in the other, and this balance of stress and relaxation is controlled by the ATP activity through phosphorylation events.

## Materials and Methods

### Chemicals and Reagents

HEK (Human Embryonic Kidney)-293T cell line was kindly provided by Dr. Shaida Andrabi (Biochemistry, University of Kashmir). Dulbecco’s modified Eagles medium (Gibco by Life Technologies), fetal bovine serum (Gibco by Life Technologies), penicillin-streptomycin solution (Gibco by Life Technologies), Tunicamycin (Tm) (Cal Biochem, USA), and Trypsin-EDTA (Gibco by Life Technologies). Antibodies used for Immunoblotting; anti-GST Antibody (Cell Signaling Technologies), IRE1α (14C10) Rabbit mAb (Cell Signaling Technologies), anti-Bid antibody (Cell Signaling Technologies), anti-IRE1 (phospho S724) antibody (Abcam), anti-XBP1 antibody (Abcam), phospho-JNK Rabbit mAb (Cell Signaling Technologies), phospho-eIF2α (Ser51) antibody (Cell Signaling Technologies), anti-ATF-4 antibody (Abcam), Bip antibody (Cell Signaling Technologies), anti-ATF-6 antibody (Abcam), anti-Grp78 antibody (Cell Signaling Technologies), anti-Mouse IgG AP-linked antibody (Cell Signaling Technologies), anti-Rabbit IgG AP-linked antibody (Cell Signaling Technologies), anti-Rabbit IgG Alkaline Phosphatase antibody (Sigma Aldrich), anti-mouse IgG Alkaline Phosphatase antibody (Sigma Aldrich), IRDye Goat anti-Rabbit IgG (LI-CORR Biosciences), IRDye Goat anti-Mouse IgG (LI-CORR Biosciences).

### Cell culture and treatments

HEK293-T cells were grown at 37°C in a humidified incubator with 5% CO_2_ in Dulbecco’s modified Eagles medium supplemented with 10% fetal bovine serum and 1 × penicillin-streptomycin solution. For overexpression studies, cells were transfected with pEBG-Bid (GST-tagged Bid) or pEBG (GST-tagged Vector) or pcDNA3.1 (+) IRE1. Cells without transfection were used as control (Mock). Cells were treated with 6μM tunicamycin for UPR induction.

### Immuno-precipitation

For immunoprecipitation, HEK293-T cells were transfected with pcDNA3.1 (+) IRE1 and pEBG-Bid / or pEBG-Bik /or pEBG-Bcl2 clones using PEI transfection reagent. Post 48hr transfection cells were treated with 6μM tunicamycin and incubated for 6 hr. As control cells transfected with pcDNA3.1 (+) IRE1 and pEBG empty vector and treated similarly. Cells without transfection were also treated with tunicamycin. Protein lysates were prepared using NETN buffer (50mM Tris-Cl, 150mM NaCl, 1% Glycerol, 1% Np-40, 0.1% SDS, 5mM EDTA, 0.5% Sodium deoxycholate, 10mM NaF, 17.5mM β-glycerophosphate, 1X PIC). 120μL of NETN buffer was added and incubated on ice for 1hr. The supernatant was collected after centrifugation at 14000rpm for 25min. Meanwhile, Protein G plus Agarose beads were washed three times with NETN buffer and incubated with 2μg of primary antibody (Anti- GST or IRE1α (14C10) Rabbit mAb) in 40μL of 1XTBS, overnight on a Nutator (rotating apparatus) at 4°C. On a consecutive day, protein lysate was added to the mixture and incubated on Nutator overnight at 4°C. On the third day of the experiment, bead solution was centrifuged at 2000rpm for 2min. Then washed thrice with NETN buffer by centrifugation. 30μL of 2X laemmili Loading Buffer was added to the samples and heated at 100ºC for 15min. Samples were then run on 12% SDS PAGE followed by western blotting using (IRE1α (14C10) Rabbit mAb or Anti- GST) antibody.

### Western Blotting

40-50μg of protein lysates were run on 10-12% SDS PAGE and transferred the PVDF membrane. The membrane was blocked in 3% blocking solution (3% BSA in 1XTBS) overnight. Blot was incubated overnight in a specific primary antibody (1:1000 to 1: 2000 dilution) at 4ºC. Secondary antibody against the mouse or rabbit primary antibody was added and incubated at room temperature for 2hr. Depending on the nature of the secondary antibody, detection assays were performed. In this study, we used two types of secondary antibodies. IRDye Goat anti-Rabbit IgG (1: 50000 dilution), Goat anti-Mouse IgG ((1: 50000 dilution) fluorophore conjugates and visualized in different fluorescence channels, 800nm and 700nm respectively with the help of Licor detection system (Odyssey). The other type of secondary antibody is an ALP conjugate (1:40,000 dilution). The blot was kept in ALP developing solution for 2-3min until bands appear on the membrane. Then the blot was washed with tap water and analyzed on the Gel doc system.

### Yeast two-hybrid assay

For yeast two-hybrid assay, Ire1 and Bid genes were cloned in pGAD424 and pGBT9 yeast vectors respectively. pGAD424 harbors GAL4 activation domain and pGBT9 contains the GAL4 transcription binding domain. Both the plasmids were co-transformed in AH109 yeast strain, which harbors a His-reporter gene. Single colonies were picked and patched on SDL^−^ T^−^ Agar and incubated at 30°C for three days. Once colonies grew, they were spotted on SDL^−^ T^−^H^−^ (SD complete media with leucine, tryptophan, and histidine drop out) plates. Plates were continuously observed for 5-6 days to assess the growth. Similarly, co-transformation of pGAD-Ire1 + pGBT9 vector and pGBT-Bid + pGAD4242 was done to set a negative control for the assay. Since the histidine biosynthesis gene promoter is responsive to the GAL4 transcription factor. Therefore, expression of His-gene will depend on the reconstitution of a GAL4 transcription factor by direct interaction between Ire1 and Bid that would allow the growth of colonies on SDL^-^T^-^H^-^ drop out media.

### siRNA knockdown

HEK293-T cell line was transfected with Mission esiRNA human BID using Lipofectamine™ 2000 reagent. Mission siRNA fluorescent universal negative control #1, Cyanine 3 was also transfected, acting as a negative control for siRNA knockdown. HEK293-T cells were seeded in five different 60mm dishes for siRNA transfection. Cells were taken for transfection at 65% confluency. Two different transfection mixtures A and B were prepared. Mixture A was prepared by mixing 1400ng of siRNA with 250μL of DMEM and incubated at room temperature for 5min. Mixture B was prepared by adding 5μL of lipofectamine to 245μL of DMEM. Mixtures A and B were mixed and incubated for 20min at room temperature. The media was removed from the dishes, and 1.5mL of DMEM was dispensed followed by the addition of transfection mixture and incubated in an incubator at 37ºC incubator. Post 12-16hr of incubation, media was removed and fresh 3mL of complete media was added. Cells were harvested at 24hr, 48hr and 72hr post siRNA transfection along with the negative control and mock (without transfection). Then samples were subjected to Western blotting to analyze the Bid knockdown by probing with Anti-Bid antibody.

### RNA isolation, Reverse transcription and real-time PCR

Total RNA was extracted from the cells using Trizol. cDNA was synthesized from total RNA with M-MLV Reverse transcriptase using random primers. 2μg of RNA was used for each reverse transcription reaction. Real-time PCR was performed with SYBRGreen fluorescent reagent to check the abundance of specific transcripts and for each reaction 50-70ng of cDNA was used. Actin or GAPDH primers were used as an endogenous control. The reaction was carried out in the 7500 Real-Time PCR System (Applied Biosystems). By normalizing the comparative threshold cycle with endogenous controls, quantitation of each transcript was done.

### Retrieving 3D structure of proteins

The crystal structures of BID protein (PDB ID: 2BID) and human IRE1protein (PDB ID: 2HZ6, PDB ID: 4U6R, PDB ID: 4YZD and PDB ID: 4Z7G) were retrieved from Protein Data Bank (http://www.rcsb.org).

### Quaternary structure analysis of IRE1

The quaternary structures analysis of IRE1 protein analysis was done using IPAC (Inference of Protein Assembly in Crystals). This program detects quaternary structures from crystal lattice by utilizing a scheme where point group symmetry works in combination with naive Bayes classifier under the Boolean framework (Mitra *et al.*, 2011). In the first step, by utilizing the asymmetric unit information available from the PDB file, we generate all symmetry-related molecules in the lattice. Non-protein atoms were also included. In this step, we also checked the presence of disulfide bonds at the interface, which was grouped as a functional unit (FUs). In the next step, biological interfaces were detected by applying Bayes classifier. Smaller interfaces were connected into a large single interface. A depth-first search was used cyclically until no new interfaces were detected. In the last step, point group symmetry (PGS) was used to screen merged FUs. Cyclic PGS is applied to identify the FUs of size n (>2). Symmetry with a threshold of 10.0 Ao can be identified by using backbone-trace Ca atoms. The largest FU satisfying PGS was inferred as the quaternary structure of the protein. In this particular study, we have used an updated version of this program that emphasizes the contribution of symmetry in determining the final quaternary structure.

### Molecular docking

Molecular docking of Bid with the cytoplasmic domain of IRE1 was performed using ZDOCK software. The docking poses were scanned using 12-degree rotations against protein 4U6R and the top 100 hits were saved. The rank 1 complex from the software output, which had a score of 808.658 was selected and taken for further investigation. The Discovery Studio Molecular Modeling client Version 4.1 facilitated the graphical visualization of the docking results.

### Pose validation

It may be noted that the Pro51-Gln81 stretch of BID is of irregular structure. The N-terminal segment Gly1 to Glu16 is also of irregular structure. Therefore, docking poses in such cases where the irregular structural segment makes a contact, they were deemed as ambiguous results and therefore discarded. Only the first ranked pose was taken for the investigation. The receptor IRE1 undergoes significant structural changes during phosphorylation-dephosphorylation events; therefore, redocking for the pose improvement was avoided.

### Statistical analysis

The results presented in this study are expressed as means ± standard error. GraphPad PRISM v6.03 statistical software (GraphPad Software, La Jolla, CA) was used for statistical analysis. One-way ANOVA was used to determine the significance of differences between different groups. A p-value of 0.05 was considered significant.

