## Supplemental Figure for "Bid as a novel interacting partner of IRE1 differentially regulating its RNAse activity"

### Supplementary Figures

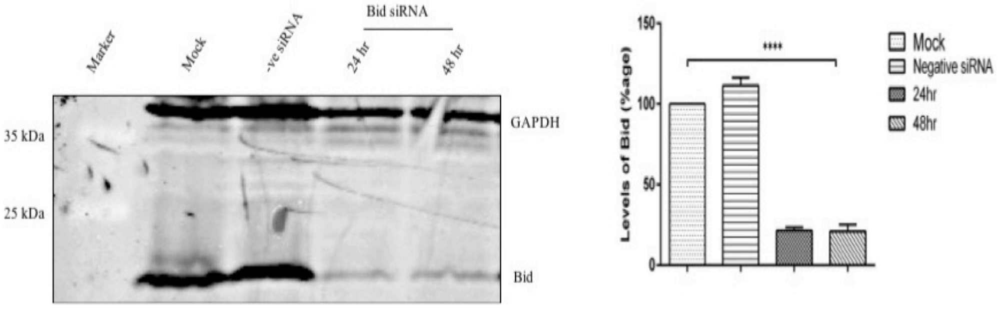


**FigureS1: Standardization of Bid siRNA knockdown.** HEK293T cells were transfected with Bid siRNA and negative control siRNA (-ve siRNA) independently. Mock was kept as a no transfection control. Cells were harvested at 24hr and 48hr post-transfection. Western blot analysis was performed from whole-cell protein lysate using with Anti- Bid antibody to detect expression levels of Bid. Densitometry quantitation of Bid levels was done using ImageJ. Here histogram represents mean SD. Statistical analysis was performed using One way ANOVA: ****: $p < 0.0001$

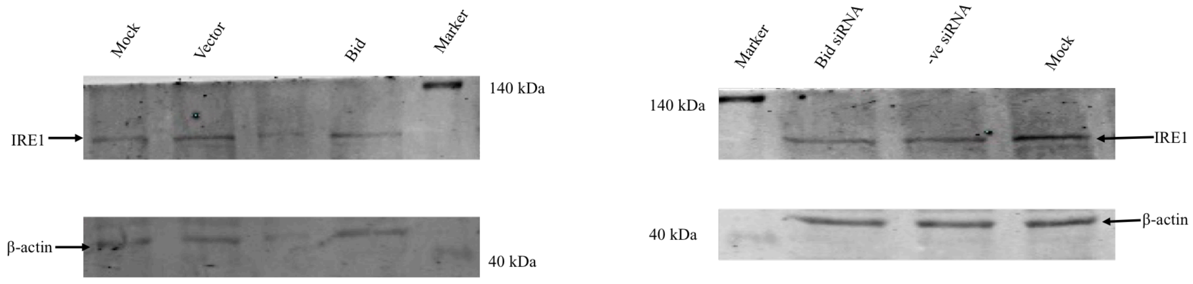


**Figure S2. Total IRE1 levels.** Analysis of the total IRE1 in Bid overexpression and knockdown conditions. HEK293T cells were transiently transfected with Gst-tagged Bid, Gst-tagged Vector and Mock (no transfection). After 42 hrs cells were stimulated with 6μM tunicamycin for 6hrs. HEK293T cells were transfected with Bid siRNA and –ve siRNA followed by treatment with 6μM tunicamycin for 6hrs. Levels of the total IRE1 were checked by probing with the anti-IRE1 antibody. β-Actin represents endogenous control. Marker represents a Pre-Stained Protein Ladder, Bid-Gst (overexpressed Bid), Vector-Gst (overexpressed vector), Mock (no transfection control), -ve siRNA (mission siRNA fluorescent universal negative control #1, Cyanine 3) and Bid siRNA (Mission esiRNA human BID).
